# Bile acids mediate liver-bone marrow crosstalk

**DOI:** 10.1101/2023.07.05.546968

**Authors:** Daniele Vitale, Mahmoud Karimi Azardaryany, Ghazal Alipour Talesh, Mahsa Shahidi, Vikki Ho, Suat Dervish, FX Himawan Haryanto Jong, Maito Suo, Jacob George, Saeed Esmaili

## Abstract

The modern dietary exposome is calorie dense and poor in nutritional quality resulting in high prevalence of fatty liver disease and an increasing incidence of cardiometabolic disease and cancer. We investigated the impact of dietary composition on the interaction between the liver and haematopoietic systems in mice. Using xenograft and chemical-induced liver cancer models, we find that liver tumours *per se* have a minimal impact on haematopoietic stem and progenitor cell (HSPC) responses. In contrast, alterations in dietary composition have profound effects on the liver-bone marrow axis. Specifically, exposure to sucrose with or without dietary cholesterol has minimal impact on the HSPC response, while perturbations in bile acid biosynthesis synergises with excess dietary cholesterol to enhance HSPC responses. Pharmacological restoration of bile acid biosynthesis partially reversed these effects. We conclude that the crosstalk between liver and bone marrow, and subsequent HSPC responses is regulated by bile acid biosynthesis.

## Introduction

The liver is a crucial metabolic hub, buffering the effects of dietary intake on the body’s internal milieu. Thus, investigating liver regulatory mechanisms can provide insights on the impact of environmental factors such as diet on the liver, and through this, the impact on other organs. While the liver’s cross-talk with other organ systems has been extensively studied, its communication with bone marrow haematopoietic stem cells remains unexplored particularly in the context of metabolic associated fatty liver disease (MAFLD) and liver cancer ^1^. We sought to understand the cross-talk between the liver and bone marrow haematopoietic stem cells for new insights on the development and progression of fatty liver and its associated diseases.

Fatty liver disease is highly prevalent; affects one in three people worldwide, and is associated with diabetes, cardiovascular disease, and cancer. Despite this, the underlying mechanisms driving both the hepatic and extrahepatic manifestations of fatty liver disease are poorly understood. We considered that understanding the gene-environment interactions that regulate metabolic health is an essential step to unravelling this complex disease ^2^. Recently, we observed an association between hepatic gene co-expression networks and bone marrow haematopoietic stem and progenitor cell (BM HSPC) responses that was triggered by dietary composition in a liver cancer model ^1^. In this model, upregulation in hepatic immune responses and downregulation in liver metabolism networks correlated with the extent of upregulation in BM HSPC responses to diet. However, this model could not distinguish whether the presence of liver cancer affects the BM HSPC response.

In this study, we explored the role of dietary factors, liver cancer, and liver metabolism on the BM HSPC response. We first show that tumours in liver cancer models have minimal effects on the HSPC response. We then demonstrate that the liver utilises bile acid biosynthesis to limit haematopoietic system activation. When bile acid synthesis is perturbed through dietary manipulation, haematopoiesis is activated and results in a heightened systemic immune response. Restoring bile acid biosynthesis using an ileal apical bile salt transport inhibitor alleviated HSPC and liver immune responses. These data suggest that by regulating bile acid metabolism we can significantly impact systemic immune responses, and that fine-tuning this process can be of therapeutic value.

## Results

### Dietary composition rather than presence of tumour regulates BM HSPC responses

Haematopoietic system changes due to release of immune modulators either by tumour cells or immune cells have been reported in cancer ^3,4^. Moreover, the common risk factors for cancer and cardiovascular diseases such as obesity and lipid abnormalities can alter haematopoiesis ^5-7^. Fatty liver disease associated with metabolic dysregulation also shares these common risk factors. In a previous study we reported the bone marrow haematopoietic responses in mice that develop fatty liver-related liver cancer while on diets of varying composition ^1^. However, that study did not allow us to distinguish whether dietary composition exerts effects on the liver and marrow independent of tumour presence, nor could we deduce if liver tumours *per se* could regulate this process.

To address the latter, we used a syngeneic tumour model by injecting Hepa1-6 liver tumour cells into the flank of immunocompetent C57BL6 mice. Tumours were harvested either 4 weeks later or when the estimated tumour weight reached 1 gram; vehicle-injected mice served as controls (Figure 1A).

**Figure 1.**
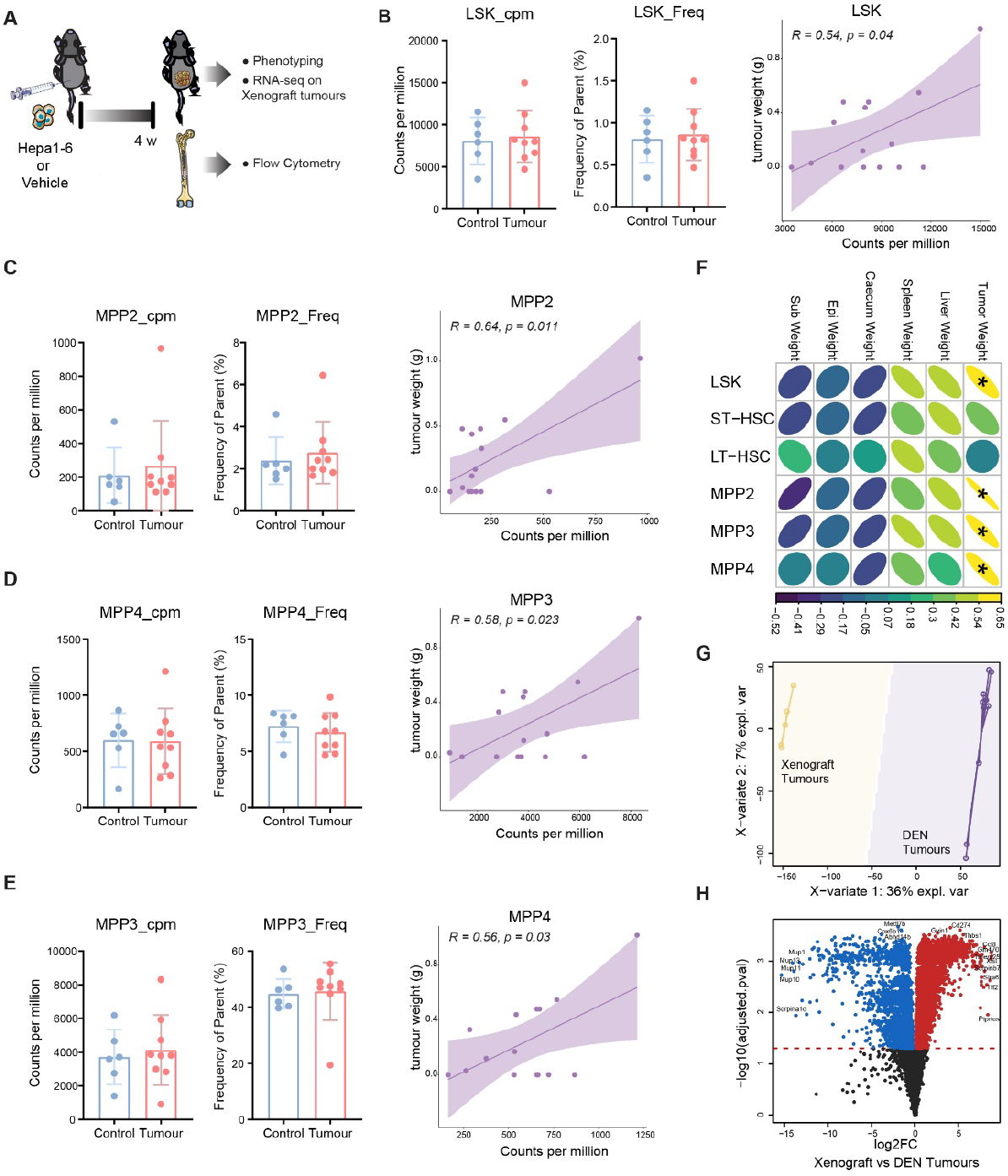
A positive association between tumour size and bone marrow HSPCs response in a xenograft tumour model. A) Schematic of xenograft tumour study with HSPCs immune profiling. B) Number and frequency of LSK (Lin-Sca1+ ckit+) cells in mice bearing tumours (n_(mice)_ = 9) and controls (n_(mice)_ = 6). LSKs number is positively correlated with tumour size (r=0.54, p=0.04). C-E) Analysis of LSK subpopulation shows a positive correlation between tumour size and the number of MPP2 (r=0.64, p=0.011); MPP3 (r=0.58, p= 0.023); and MPP4 (R= 0.56, p=0.03). F) Pairwise comparisons arranged in a matrix where the direction and magnitude of Pearson’s correlation between phenotypic variables is represented by the colour of an ellipse whose pitch and eccentricity inform the direction and magnitude, respectively. Statistically significant correlations are marked. G) PLS-DA analysis of RNA-seq in xenograft tumours (n_(mice)_ = 5) and DEN (n_(mice)_ = 10) induced tumours. Xenograft tumours are separated from DEN tumours. H) Volcano plot shows large number of differentially expressed genes between xenograft and DEN induced tumours fed the NC diet.

Flow cytometry-based immunophenotyping of bone marrow haematopoietic cells revealed no differences in the number and frequencies of lineage-negative (Lin)- Sca-1^+^ c-Kit^+^ (LSK) cells, haematopoietic stem cells (HSCs), or multipotent progenitor cells 2, 3, and 4 (MPP2, MPP3, and MPP4) between tumour-bearing mice and controls (HSCs and MPPs are collectively called hematopoietic stem and progenitor cells). However, we observed a positive correlation between tumour size and the number of LSK, MPP2, MPP3, and MPP4 cells (Figure 1B-F). This suggests that very large tumours might regulate HSPC responses, either due to the presence of tumour or because the animals are stressed in the presence of large tumours, as previously demonstrated ^8^.

To investigate the differences between the tumour models, we performed RNA-seq on xenograft tumours and compared them with tumours induced previously using diethylnitrosamine (DEN). Partial least squares discriminant analysis (PLS-DA) and volcano plots of differentially expressed genes (DE genes) revealed distinct gene expression profiles between xenograft and DEN-induced tumours (Figure 1G-H). Given the differences in gene expression profiles between these models, we focused our subsequent studies on the DEN-induced tumour model.

To investigate the effects of diet composition on DEN-induced liver tumorigenesis, we subjected 22 week old mice to 14 weeks of diets that varied in their composition. We used high sucrose (HS), normal chow with cholic acid 0.5% (CA), or HS_Chol2%_CA (a cholesterol rich diet with cholic acid) diet challenges, while control mice were DEN-injected and fed normal chow (NC) diet (Figure 2A). Mice exposed to the HS diet had higher body weights and adipose tissue mass and exhibited a trend towards higher blood glucose levels (Figure 2B-D). Mice fed the HS_Chol2%_CA diet had higher blood cholesterol levels (Figure 2E). Both CA and HS_Chol2%_CA diets increased liver weight and liver/body weight ratio (Figure 2F). Plasma ALT levels were elevated in the HS, CA (p<0.05), and in the HS_Chol2%_CA diet compared to control mice (Figure 2G). These findings suggest that dietary composition differentially impact metabolic and liver-related parameters in the context of DEN-induced tumorigenesis.

**Figure 2.**
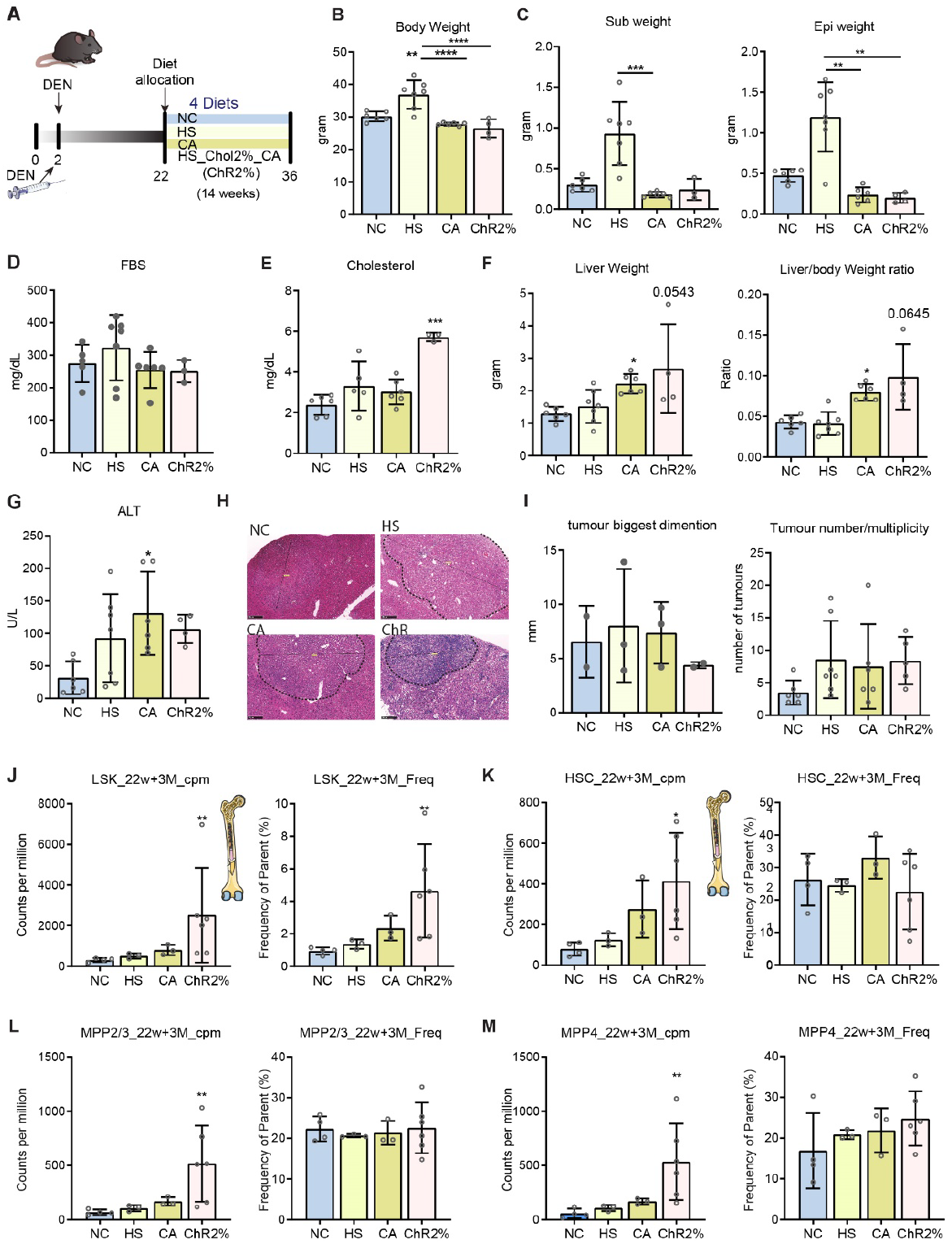
Short-term dietary challenge in a DEN tumour model did not show a synergism between diet and DEN tumours. A) Schematic shows the experimental plan with DEN injection (25 mg/kg) at the age of 2 weeks, 14 weeks of dietary challenge starting at 22 weeks of age (n= 5-7 mice/group). B-C) HS diet increased body and adipose tissue weights. D) A higher trend in fasting blood glucose in HS diet fed mice. E) Plasma cholesterol was highest in ChR2% diet fed mice. F) Liver weight and liver/body weight ratio was higher in CA and ChR2% diet fed mice. G) Higher ALT levels in HS (a trend), CA (p<0.05), and ChR2% (a trend) diet fed mice. H) Representative of liver H&E staining in diet groups. I) Similar tumour burden between ChR2% and NC diet fed mice. J) Number and frequency of Lin-Sca-1+ cKit+ cells (LSKs) increased in ChR2% diet group, with a higher trend in CA diet fed mice. K) The number and frequencies of HSCs (long-term and short-term HSCs) between diet groups. L-M) Higher trends in number and frequency of multipotent progenitor cells 2/3 (MMP2/3), and MPP4 in ChR2% diet group. Error bars represent mean ±1 standard deviation (SD). One-way ANOVA was used to test for significant differences in means with Tukey’s (parametric), and Dunn’s (non-parametric) multiple comparison post-hoc test. ^****, ***, **^ and ^*^ indicates a significant difference compared to normal chow, with P <0.0001, P<0.001, P<0.01 and P<0.05 respectively. (NC, normal chow, n_(mice)_ = 6; HS, high sucrose, n_(mice)_ = 7; CA: cholic acid, n_(mice)_ = 6 HS_Chol2%_CA, high sucrose + high cholesterol (2%) + cholic acid, n_(mice)_ = 6; a fraction of the samples was used in some of the experiments)

We did not detect any significant difference in the liver tumour burden between the mice groups exposed to these dietary interventions (Figure 2H-I). Importantly, flow cytometry of HSPCs from the mice showed a lack of response to the HS diet (compared to NC) and a trend to an elevated HSPC response in mice fed the CA diet. In contrast, the HS_Chol2%_CA diet group showed significantly higher BM HSPCs responses (Figure 2J), with no differences observed in their subpopulations (Figure 2K-M). In sum, these findings suggest that DEN-induced tumours do not impact the HSPC response on the NC or HS diets while a diet comprising sucrose, cholesterol, and cholic acid shows significant activation of the HSPC response.

### The role of bile acids in regulating bone marrow HSPC responses

To further investigate the interplay between liver and bone marrow, we utilized a series of non-tumour dietary models to induce an array of liver pathologies. The diets encompassed different combinations of sucrose and cholesterol, with and without cholic acid (CA), namely: A) normal chow (NC); B) high sucrose (HS); C) high sucrose + high cholesterol (2%) (HS_Chol2%); D) high sucrose + high cholesterol 2% + cholic acid 0.5% (HS_Chol2%_CA); E) high cholesterol 2% + cholic acid 0.5% (Chol2%_CA); and F) cholic acid 0.5% (CA) (Figure 3A).

**Figure 3.**
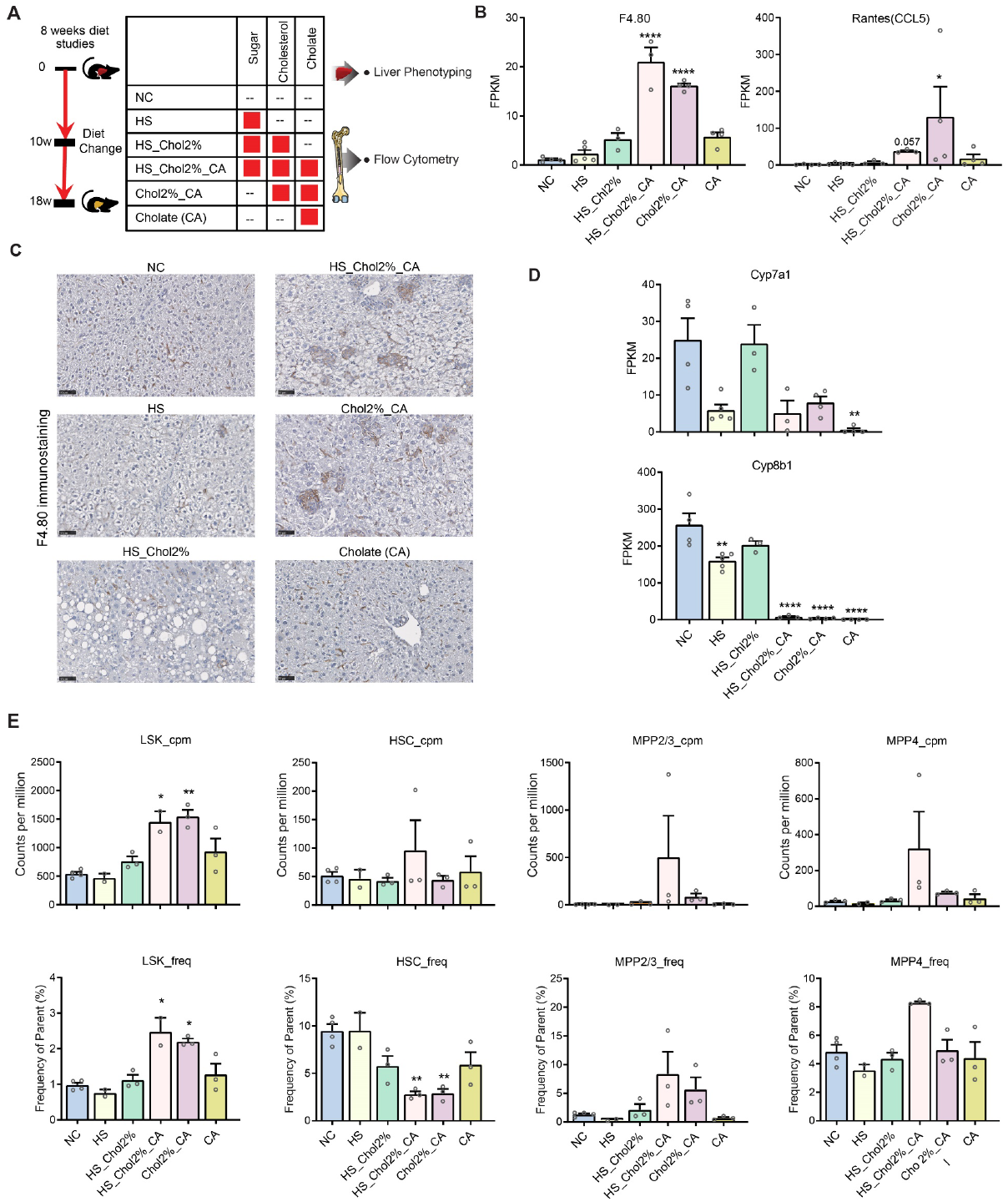
Dietary challenge in mice induced a spectrum of liver pathologies and bone marrow haematopoiesis responses. A) Schematics show the composition of 6 diets. B) Liver inflammatory markers are upregulated in diets containing cholesterol and cholic acid. C) Immunostaining of F4.80 shows high presence of macrophages in the livers of mice fed the diets containing cholesterol and cholic acid. D) Suppression of bile acid biosynthesis in mice expose to cholic acid. E) Higher number of LSK in mice fed the diet containing cholesterol and cholic acid. (3-5 mice/group were used. In flow cytometry, outliers identified using Grubbs test.)

The diets were designed to elicit varying degrees of liver pathologies over an 8-week period ^9^. The expression of the macrophage marker F4/80 and the inflammatory marker Rantes within the liver was upregulated in diets that contained both CA and cholesterol: HS_Chol2%_CA, and Chol2%_CA (Figure 3B). Notably, no significant difference was observed between HS_Chol2%_CA, and Chol2%_CA diets, indicating a synergistic effect of CA with cholesterol. We noticed a higher trend in expression of macrophage markers in the HS_Chol2%, and CA diet fed mice (Figure 3B). Immunostaining for F4.8 confirmed significant accumulation of macrophages in diets containing both cholesterol and CA (Figure 3C). The HS_Chol2% diet showed some degree of immune infiltration, while the high sucrose diet alone failed to induce inflammation. Examining liver metabolic response to dietary challenge showed that the expression of genes related to bile acid synthesis such as *Cyp7a1* and *Cyp8b1* was consistently and significantly lower in mice exposed to diets with added CA (Figure 3D). Exposure to sucrose also reduced expression of these markers (Figure 3D).

Investigation of bone marrow haematopoietic stem cells via flow cytometry analysis of mice on these diets revealed an elevated number and percentage of Lin-Sca1+ ckit+ (LSK) and haematopoietic stem and progenitor cells (HSPCs) in diets enriched in cholesterol *and* cholic acid (Figure 3E). However, the frequency of HSCs was reduced in these diets. Importantly, the HS_Chol2% diet (without CA) and CA diet did not have a significant impact on the BM-HSPCs response (Figure 3E).

Further analysis of the HSPC sub-population indicated a significant increase in the population of multipotent progenitor cells (MPP2, MPP3, MPP4) in diets enriched in cholesterol *and* cholic acid. The MPP2/3 are myeloid biased cells and generate all myeloid linages, while MPP4 cells are biased toward lymphoid cells ^10,11^. The number of HSCs (short-term and long-term HSCs) was not different between diets, although their frequencies were decreased in diets that contained cholesterol and cholic acid (Figure 3E). This decrease in HSC frequency can be attributed to the fact that the majority of LSKs (the parent cells utilised to calculate HSC frequencies) in diets that contained cholesterol and cholic acid were of the multipotent progenitor cell population. Our findings suggest a systemic response and overall activation of BM HSCs as well as their differentiation into progenitor cells in diets that contain both cholesterol and cholic acid (Figure 3E).

The above data suggest a critical role for cholic acid in activating bone marrow hematopoietic stem and progenitor cells (HSPCs) in the context of a high cholesterol diet. To investigate the possibility of reversing the effects of added cholic acid, we used SC-435 (Figure 4A), a selective inhibitor of the apical sodium-dependent bile acid transporter (ABST) to reduce bile acid absorption in the gut ^12^. Treatment with SC-435 reduced liver injury and inflammation in the HS_Chol2%_CA diet model (Figure 4B). Notably, SC435 treatment also resulted in a significant reduction in liver cholesterol levels (P<0.01), yet it remained far above normal levels. (Figure 4C). Moreover, SC-435 treatment enhanced the mRNA expression of Cyp7a1, a marker of liver bile acid synthesis in the NC group. In mice on the HS_Chol2%_CA diet, Cyp7a1 mRNA expression was suppressed, but SC-435 treatment reversed this inhibitory effect of cholic acid (Figure 4D). Further analysis of bone marrow flow cytometry data revealed a consistent reduction in the number and frequencies of LSK, and HSCs in HS_Chol2%_CA diet-fed mice treated with SC-435 (Figure 4E-F). Taken together, our findings suggest that high dietary cholesterol and dysregulated bile acid biosynthesis have a synergistic multi-system effect that significantly impact the BM HSPC response.

**Figure 4.**
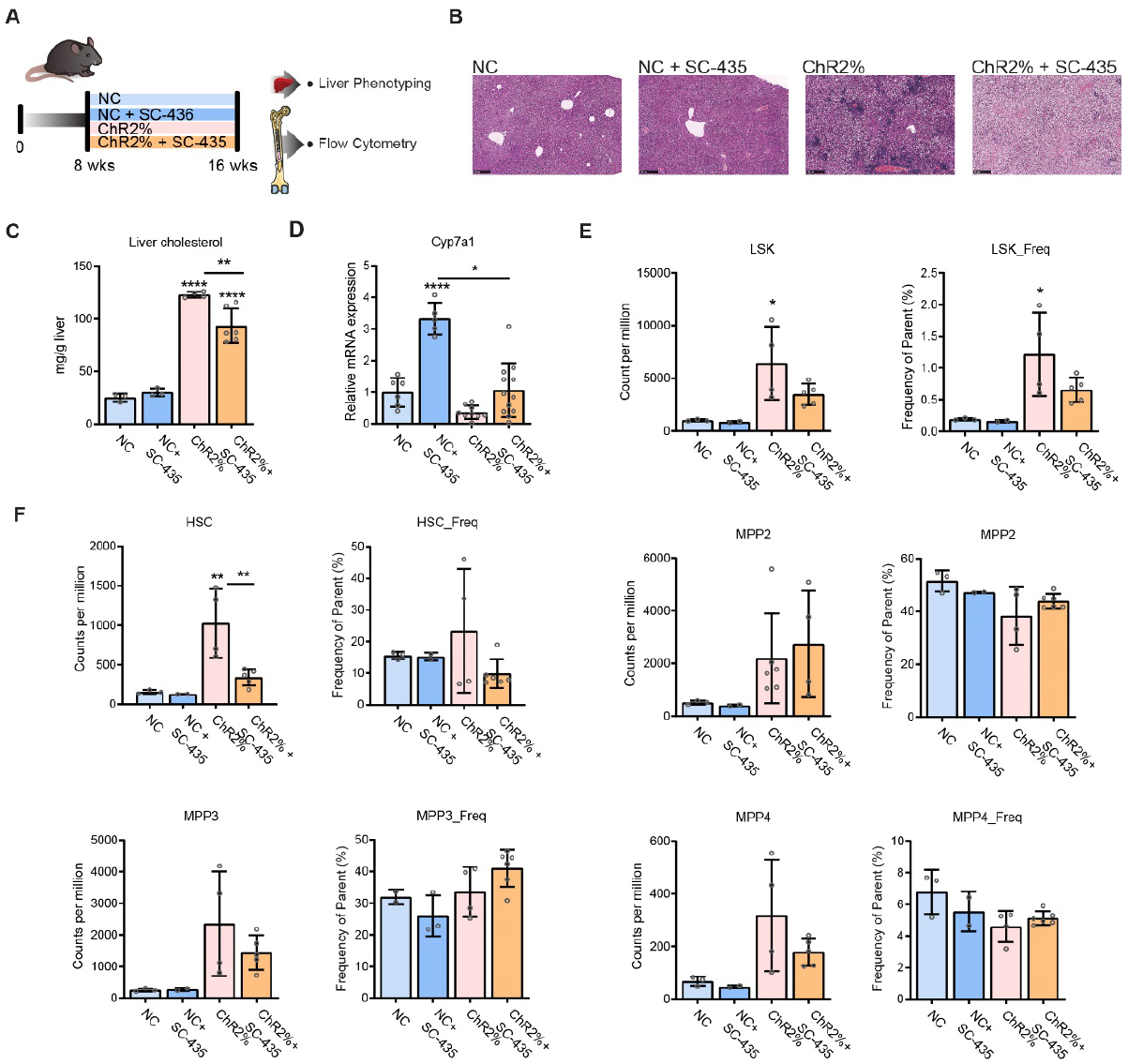
Regulating bile acids metabolism can regulate BM HSPCs response. A) Schematics of an interventional study using HS_Chol2%_CA diet with/out SC-435. B) Representative H&E staining shows SC-435 treatment alleviated liver immune cells infiltration. C) Mice fed the ChR2% diet showed increased total liver cholesterol while adding SC-435 slightly reduced total liver cholesterol. D) SC-435 increased liver Cyp7a1 mRNA expression. E) Feeding mice the ChR2% diet significantly increased the number and frequency of LSK cells, while SC-435 markedly alleviated LSKs response. F) Analysis of LSK subpopulation shows reduction in the number of HSCs with lower trend in MPP3 and MPP4. (NC, normal chow, n_(mice)_ = 6; NC +SC-435, n_(mice)_ = 5; HS_Chol2%_CA (ChR2%), high sucrose + high cholesterol (2%) + cholic acid, n_(mice)_ = 10; ChR2% + SC-435, n_(mice)_ = 13; 3-5 mice/group were used in flow cytometry, outliers identified using Grubbs test.) Error bars represent mean ±1 standard deviation (SD). One-way and two-way t-test was used to test for significant differences in means with Tukey’s (parametric) and Dunn’s (non-parametric) multiple comparison post-hoc test. ^****, ***, **^ and ^*^ indicates a significant difference compared to normal chow, with P <0.0001, P<0.001, P<0.01 and P<0.05 respectively.

## Discussion

The interaction between the liver and bone marrow hematopoietic system is relatively underexplored. To address this knowledge gap, we investigated how the dietary exposome affects liver regulatory mechanisms that control the hematopoietic stem and progenitor cell (HSPC) response. We found that the presence of liver tumours in two models was not associated with any alteration in the HSPC response, Dietary composition on the other hand played a crucial role. Specifically, exposure to sucrose and cholesterol, as well as liver cholesterol accumulation had minimal impact on BM HSPCs. However, excess dietary cholesterol together with the perturbation of bile acid biosynthesis (with the addition of cholic acid) synergistically induced an HSPC response. Importantly, rescuing bile acid synthesis alleviated the HSPC response. These findings suggest that liver homeostatic dysregulation, along with bone marrow overstimulation, could underly at least some of the extra-hepatic manifestations of fatty liver disease.

Exposure to cholesterol in conjunction with disrupted bile acid homeostasis via cholic acid had a significant impact on hematopoietic stem and progenitor cells responses. Notably, restoring bile acid synthesis using a pharmacological intervention effectively alleviated the bone marrow HSPCs response, despite the presence of high levels of liver cholesterol. Thus, dietary cholesterol exhibited synergistic effects with added cholic acid in the diet and resulted in the induction of an HSPCs response. Though the combination of high sucrose diet with added dietary cholesterol induced liver inflammation, it had minimal impact on HSPC responses. In sum, these findings highlight the importance of liver bile acid metabolism on the HSPCs response.

It is important to acknowledge the limitations of model systems in studying human diseases and how best to extrapolate such findings to humans. We showed that mere dysregulation in cholesterol homeostasis was not sufficient to incite BM HSPC responses in mice, while suppression of bile acid biosynthesis was. Of note, we have previously shown the protective effects of inhibiting bile acid and cholesterol biosynthesis on reducing liver cancer burden ^1^. Interestingly, a recent analysis of patients in the IMBrave 150 clinical trial (immunotherapy in HCC) showed that responders had down-regulation in pathways related to cholesterol homeostasis and bile acid synthesis ^13^. Conceivably, targeting these pathways (e.g., using statins or FXR agonists such as Obtecolic acid) could serve as adjuvants for immunotherapy response in liver cancer.

In conclusion, we demonstrate that the dietary exposome impacts liver as well as systemic homeostasis, at least at the level of the haematopoietic stem cell compartment. We show that the systemic effect is modulated, at least in part, by the regulation of bile acid production. Thus, controlled manipulation of liver-bone marrow interactions might be a promising avenue for developing novel therapeutics for both the hepatic and extra-hepatic complications of fatty liver disease.

## Acknowledgements

Flow cytometry was performed at the Westmead Scientific Platforms, which are supported by the Westmead Research Hub, the Cancer Institute New South Wales, the National Health and Medical Research Council and the Ian Potter Foundation. S.E., and J.G. are supported by the Robert W. Storr Bequest to the Sydney Medical Foundation, University of Sydney; a National Health and Medical Research Council of Australia (NHMRC) Program Grant (1053206) and Project grants (APP1107178 and APP1108422). Mice tumour studies was supported by Sydney Medical School Foundation. G.A.T. and D.V. are supported by a scholarship from the University of Sydney.

## MATERIALS AND METHODS

### Mice and diet

All procedures were approved by the Western Sydney Local Health District Animal Ethics Committee and conducted in accordance with Animal Experimentation guidelines of the National Health and Medical Research Council (NHMRC) of Australia. For the DEN model study, male C57BL6 mice were injected intraperitoneally with 25 mg/kg body weight DEN (Sigma-Aldrich, Munich, Germany) at 14 days of age. These mice were then given the diets containing 34% Sucrose (HS diet), cholic acid (0.5%) added diet (CA), or a cholesterol rich diet (ChR 2%) which contained high sucrose (34%), 2% cholesterol and 0.5% cholic acid (Specialty Feed Service, Glen Forest, Australia) starting from week 22 for 14 weeks. The control group was fed with the NC diet throughout the experiment. The mice were harvested at 34 weeks after injection (at the age of 36 weeks). Mice were exposed to a 12-hr light/ dark cycle in pathogen free conditions with free access to food and water. In non-DEN diet studies, mice were given the NC, HS, HS_Chol2%, HS_Chol2%_CA, Chol2%_CA, or CA (0.5%) diets for 8 weeks starting at 8-10 weeks of age. In pharmacologic interventional experiments, mice were fed with NC or HS_Chol2%_CA (ChR2%) diets with/out SC-435 for 8 weeks starting at 8-10 weeks of age, as we reported previously^12^.

### Biochemical assays

Serum ALT and AST were measured by automated techniques within the Department of Clinical Chemistry, Westmead Hospital. Serum lipid panel was measured using Cobas b 101 POC system and its lipid disc (both from Roche Diagnostics). For liver cholesterol measurements, total lipids were extracted using the method described by Shimabukuro et al ^14^. Briefly, 30mg of frozen liver was used to extract total lipids with 3:2 tert-butyl alcohol:triton X-100/methyl alcohol (1:1). Liver cholesterol levels were determined by enzymatic colorimetric assays using Wako E Total Cholesterol Kit (Cat # 439-17501) according to the manufacturer’s protocol.

### Immunostaining

Heat-induced epitope retrieval was applied on formalin fixed paraffin embedded (FFPE) liver tissue sections (4 μm) with 1x Rodent Decloaker buffer (Biocare Medical Inc., Concord, CA, USA) at 95°C for 40 minutes. Sections were incubated with peroxidase blocking solution Bloxall) for 10 minutes. After washing in TBST, slides were incubated with DAKO Protein Block, serum-Free (DAKO, X0909) for 45 minutes, followed by overnight incubation with F4/80 antibody (Clone CI:A3-1, Bio-Rad Laboratories) diluted in DAKO antibody diluent (1:100) with background reducing components (DAKO, S3022). For DAB staining, A biotin labelled secondary Ab (goat anti rabbit, #31820 Invitrogen), 1:500 diluted in DAKO antibody diluent was incubated for 1 hour at RT. Subsequently, slides were washed in TBST and incubated with Streptavidin HRP (SA-5004 Vector Laboratories) for 10 min at RT. ImmPACT DAB Substrate (SK-4105 Vector Laboratories) was used for developing and counterstained with Mayers Hematoxylin for 1 minute.

### Bone marrow cell isolation

Femurs were removed from mice using forceps and surgical scissors, cleaned of muscle and connective tissue and placed in wells of 6-well plates containing 3ml of cold PBS. 0.5ml tubes were pierced using a 21G syringe and inserted into 1.5mL tubes. Bone ends were cut using a scalpel and centrifuged by fast ramp up to 12,000g to extract the bone marrow. All centrifugations were done at 4°C. Bone marrow pellets were resuspended in 400ul of IMDM + 2% FBS then passed through 70um strainer, flushing through with 10ml of the medium into a 50ml falcon. Cells were pelleted at 300g for 5 mins and RBC lysed using Ammonium Chloride Lysis Buffer (Stem Cell Technologies) as per manufacturer protocol. Cells were washed twice in IMDM + 2% FBS and kept on ice.

### Flow cytometry

Prepared cell pellets were suspended in 1000 μl FACS buffer (PBS containing 5% FBS (heat inactivated) and 5mM EDTA). The live singlets were counted using DAPI 1:10 dilution by mixing 45 μl of cell suspension with 5 μl DAPI by gating on FSC-A vs FSC-H measuring average event/ μl on Cytoflex (Beckman Coulter). Subsequently, 10 million bone marrow cells/sample were transferred to round bottom 96 well plates and pelleted by spinning at 500g for 5 mins. Cell viability assessed by incubation with 50μl Zombie Yellow (BioLegend Cat. 423103) viability stain (1:500 in PBS) for 15 min in a dark, at room temperature. The cells were then topped up with 200 μl PBS, spun down at 500 x g for 5 min at 4°C. The supernatant was gently aspirated, and the pellet suspended in 100 μl of antibody cocktail in Brilliant violet (BV) stain buffer for 30 min in the dark, at room temperature. The cells were topped with 200 μl FACS buffer, spun down at 500 x g for 5 min at 4°C. The supernatant was then gently aspirated, and the pellet was washed 3 times with 200 μl FAC buffer and spun down at 500 x g for 5 min at 4°C. After the final wash, the pellet was suspended in 300 μl FACS buffer and transferred to a FACS tube and kept on ice in the dark. Fluorometric data were then acquired on a flow cytometer (Fortessa, Becton Dickinson). One well of unstained cells was included in all experiments. Single colour controls (SCCs) were prepared by adding only one single antibody to the cells in FACS buffer. Fluorescence minus one (FMO) controls were prepared by adding all the antibodies of each panel except for one to the FACS buffer. A pooled mixture of all samples was used for unstained wells, SCC and FMOs.

We used the following antibodies for flow cytometry: CD117 APC 2B8 (BD, Cat. 553356), CD140A PE-CF594 APA5 (BD, 562775), CD16/CD32 BUV737 2.4G2 (BD, 565272), Brilliant Violet 510 anti-mouse CD127 (IL-7Ra) (Biolegend, Cat. 135033), Brilliant Violet 650 anti-mouse CD150 (SLAM) (Biolegend, Cat. 115932), Brilliant Violet 785 anti-mouse CD90.2 (Biolegend, Cat. 105331), Brilliant Violet 711 anti-mouse Ly-6A/E (Sca-1) (Biolegend, Cat. 108131), APC/Cy7 anti-mouse CD48 (Biolegend, Cat. 103432), Anti mouse CD34 BV421 (BD, Cat. 562608), Anti mouse CD45 BUV395 (BD, Cat. 564279), FITC Anti mouse/rat CD29 (Biolegend, Cat. 102205), PE/Cy7 anti-mouse CD105 (Biolegend, Cat. 120410), Ms CD135 PE A2D10.1 (BD, Cat. 553842), Brilliant Violet 650™ anti-mouse/human CD44 Antibody (Biolegend, Cat. 103049).

### Analysis of liver tumours, histology and immunohistochemistry

Liver histology was assessed on 5μm paraffin embedded sections stained with haematoxylin and eosin (H&E).

### Gene expression analysis

The extraction of total RNA from liver or tumour tissues were performed with a Favor Prep Total RNA Mini Kit (Favorgen Cat. FABRK 000, FABRK 001) according to the manufacturer’s instructions. RNA concentration was measured using Nanodrop spectrophotometer (Thermo Fisher). Gene expression was measured by qPCR or by RNA-seq for tumour xenografts, as we reported recently ^1^.

### Statistical analysis

Data are presented as mean ±1 standard deviation (SD and analysed using Graphpad Prism (Graphpad). The two-tailed student t-test (Gaussian distribution), or Mann-whitney test were used for comparison of two groups. One-way analysis of variance (ANOVA) with Tukey’s (parametric) or Dunn’s (non-parametric) method for multiple comparisons were used for comparison of several groups. We did not use a statistical test to predetermine the sample size. We used a randomisation method for diet allocation. Statistical significance is presented as P<0.05 (*), P <0.01 (**), P<0.001 (***), or P<0.0001 (****).

## Notes

**Conflicts of interest:** All authors disclose no conflicts.

### Competing Interest Statement

The authors have declared no competing interest.

